# Zebrafish embryos are more efficient and robust than zebrafish larvae in evaluating human norovirus infectivity

**DOI:** 10.1101/2022.11.28.518296

**Authors:** Malcolm Turk Hsern Tan, Zhiyuan Gong, Dan Li

**Affiliations:** Department of Food Science & Technology, Faculty of Science, National University of Singapore, Singapore; Department of Biological Sciences, Faculty of Science, National University of Singapore, Singapore

**Keywords:** Human norovirus, infectivity, zebrafish embryo, UV

## Abstract

This study reports an essential improvement of the method for replication of human norovirus (hNoV) with the use of zebrafish (*Danio rerio*) embryos. With three globally prevalent hNoV genotypes and P-types GII.2[P16], GII.4[P16], and GII.17[P31], we demonstrated that this tool had high efficiency and robustness, and enabled continuous virus passaging. This tool is versatile in being applied in hNoV related research. In pathogenesis study, the zebrafish embryo generated hNoVs showed clear binding patterns to human histo-blood group antigens (HBGAs) in human saliva by a simple saliva-binding reverse transcription-quantitative polymerase chain reaction (RT-qPCR). In disinfection study, it was shown that a dose of 6 mJ/cm^2^ UV_254_ was able induce > 2-log reduction in hNoV infectivity for all three hNoV strains tested, suggesting that hNoVs were more UV susceptible than multiple enteric viruses and commonly used hNoV surrogates as tested before.

**IMPORTANCE:** HNoVs are a leading cause of gastroenteritis outbreaks worldwide. The zebrafish embryo tool as developed in this study serves as an efficient way to generate viruses with high titers and clean background and a straightforward platform to evaluate hNoV inactivation efficacies. It is expected that this tool will not only benefit epidemiological research of hNoV but also be used to generate hNoV inactivation parameters which are highly needed by the water treatment and food industry.

## INTRODUCTION

In recent years, human noroviruses (hNoVs) continue to be recognized as a leading cause of acute gastroenteritis outbreaks worldwide (1). In the United States, a total yearly cost of $10.6 billion was estimated as the economic burden of hNoV gastroenteritis (2). In China, 556 hNoV outbreaks were recorded during the period from October 2016 to September 2018 (3). Following the COVID-19 containment measures, multiple countries including Germany, Australia, and US have reported decrease in hNoV infections (4–6). However, it has been predicted that hNoV infection would resurge soon due to the relaxation of nonpharmaceutical interventions associated with COVID-19 restrictions (7).

HNoVs are extremely contagious and the transmission of hNoVs can happen via direct person-to-person contact or indirectly through fecal or vomit contaminated foods, water, fomites and air (8, 9). Therefore, researchers have dedicated numerous amount of efforts in evaluating the efficacies of inactivation strategies on the viruses (10–12). In order to do so, one must concur the challenge of hNoV cultivation and viability determination, which remained impossible for a long time in the near history (13) and is still more difficult than most of the laboratory cultivable viruses (14–17). For instance, the most recently optimized human intestinal enteroids (HIEs) model in support of hNoV replication is in need of human biopsy specimens obtained from surgery or endoscopy and five different media to maintain and differentiate (15). In comparison, the zebrafish (*Danio rerio*) larvae platform is more widely available at universities/research centers and is more cost effective for high-throughput studies (16–19). However, high individual variances of virus levels detected from the zebrafish larvae were noticed even at the peak period of virus replication (16–19). Here, we report an improvement of this model using zebrafish embryos instead of zebrafish larvae resulting significantly higher efficiency and robustness.

## RESULTS AND DISCUSSION

### Zebrafish embryos are more efficient and robust than zebrafish larvae in supporting hNoV replication and enabled continuous passaging

The clinical (stool) sample of hNoV GII.4[P16] (8 log genome copies/g stool) was tested by injecting 3 nL of the virus suspension in the yolk of each larva at 3 days post-fertilization (dpf) (same procedure as described previously [18, 19]) and in the yolk of each embryo at 6 hours post-fertilization (hpf). Using phenol red to mock the virus suspension, it was shown that the manipulation of embryo yolk injection was much easier with less resistance and thus more time-efficient than the larva yolk injection as shown in the videos (supplementary material 1 and 2).

At each time point (0 day post-infection [dpi] was taken as immediately after injection) as shown in Figure 1, 10 injected embryos or larvae were pooled as one sample and proceeded to homogenization, RNA extraction and reverse transcription-quantitative polymerase chain reaction (RT-qPCR) for hNoV GII detection. Being consistent with our previous reports (18, 19), when the viruses were injected to the zebrafish larvae (Figure 1a), low levels of hNoV below 2 log genome copies/10 larvae were recovered at 0 dpi, and a significant increase of virus genome copies (4.4 ± 0.8 log genome copies/10 larvae) were detected at 2 dpi (P < 0.05). The viral loads as detected from the zebrafish larvae started to decrease gradually from 3 dpi. At 5 dpi, the viruses already dropped to levels with no significant difference in comparison with 0 dpi (P > 0.05).

**Figure 1.**
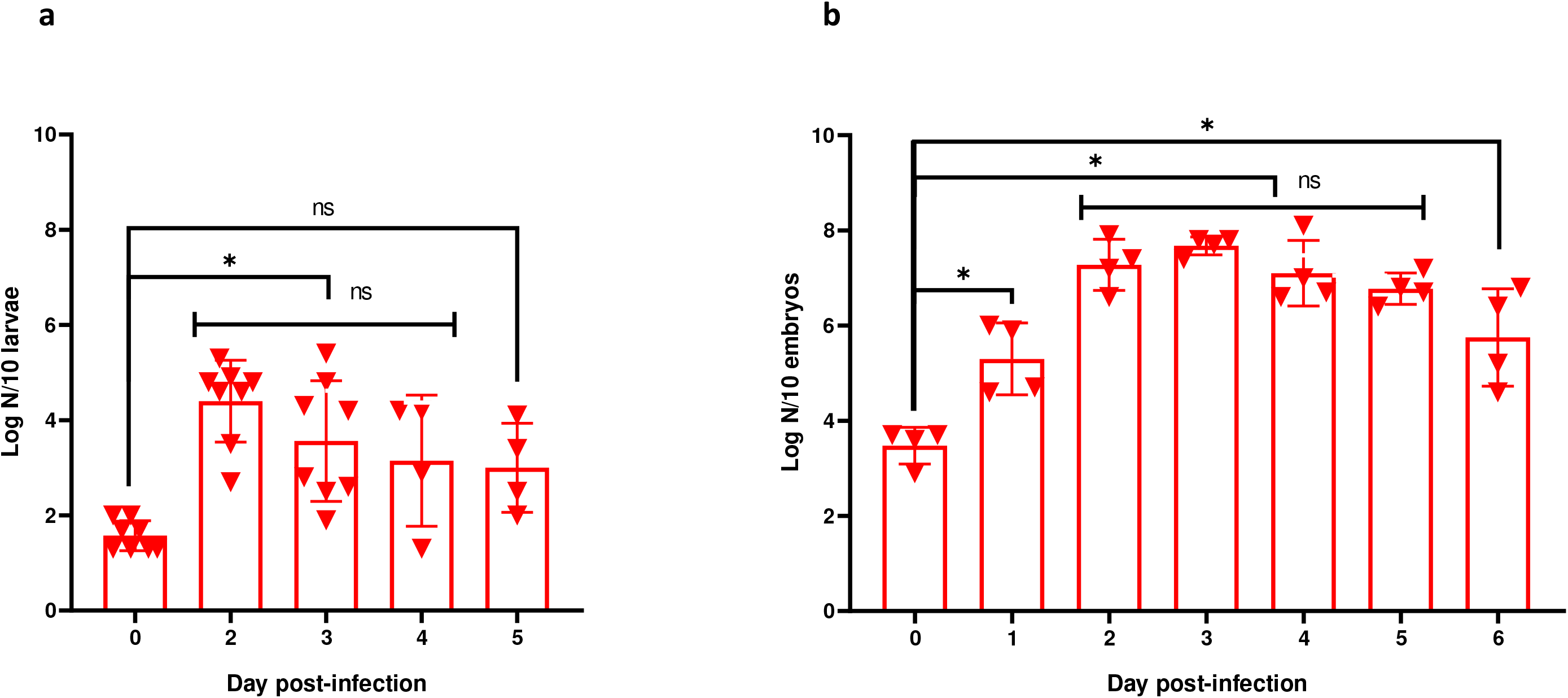
RT-qPCR detection of hNoV GII from zebrafish larvae (a) and embryos (b) at different time point. Each data point was from a pool of 10 larvae or embryos. Each column represents the average the independent biological samples, and each error bar indicates the standard deviations. *, significantly different with P < 0.05; ns, no significance with P > 0.05.

In comparison, when the viruses were injected to the zebrafish embryos (Figure 1b), higher levels of hNoV above 3 log genome copies/10 embryos were recovered at 0 dpi. This was possibly due to less virus loss during the injection of embryos than larvae, which were with more rigid structure to prick (supplementary material 1 and 2). Significant virus replication was already noticed from 1 dpi and lasted to 6 dpi which was the last day as tested (P < 0.05). The peak levels of virus detection reached at 2 dpi and lasted for 4 days (no significance from 2 dpi to 5 dpi [P > 0.05]). The virus levels detected at 3 dpi were with the smallest variation (7.7 ± 0.2 log genome copies/10 embryos) and thus 3 dpi (4.2 ± 0.4 log genome copies/10 embryos increase in comparison with 0 dpi) was selected as the time point to evaluate the virus infectivity in the following studies.

HNoVs are genetically diverse RNA viruses with 10 genogroups (GI-GX) and 49 genotypes (20). Next to GII.4[P16], we have also tested two other strains GII.2[P16] and GII.17[P31] in this study. All three strains belong to the globally prevalent genotypes and/or P-types in recently years (21). When the same protocol was applied to hNoV GII.2[P16] and GII.17[P31] (both with ~ 8 log genome copies/g stool), comparable results with hNoV GII.4[P16] were obtained at 3 dpi (P > 0.05) (Figure 2). In addition, this high level of virus replication enabled continuous passaging for all three tested strains up to four passages as tested (Figure 2). Significant increase of virus titers were obtained for hNoV GII.4[P16] (from passage 4, P < 0.05) and GII.17[P31] (from passage 3, P < 0.05). For hNoV GII.2[P16], although no significance was obtained (P > 0.05), a trend of increase in virus titers along with the passaging was also observed (Figure 2). In comparison, the zebrafish larvae only allowed successful passaging up to the second passage (16).

**Figure 2.**
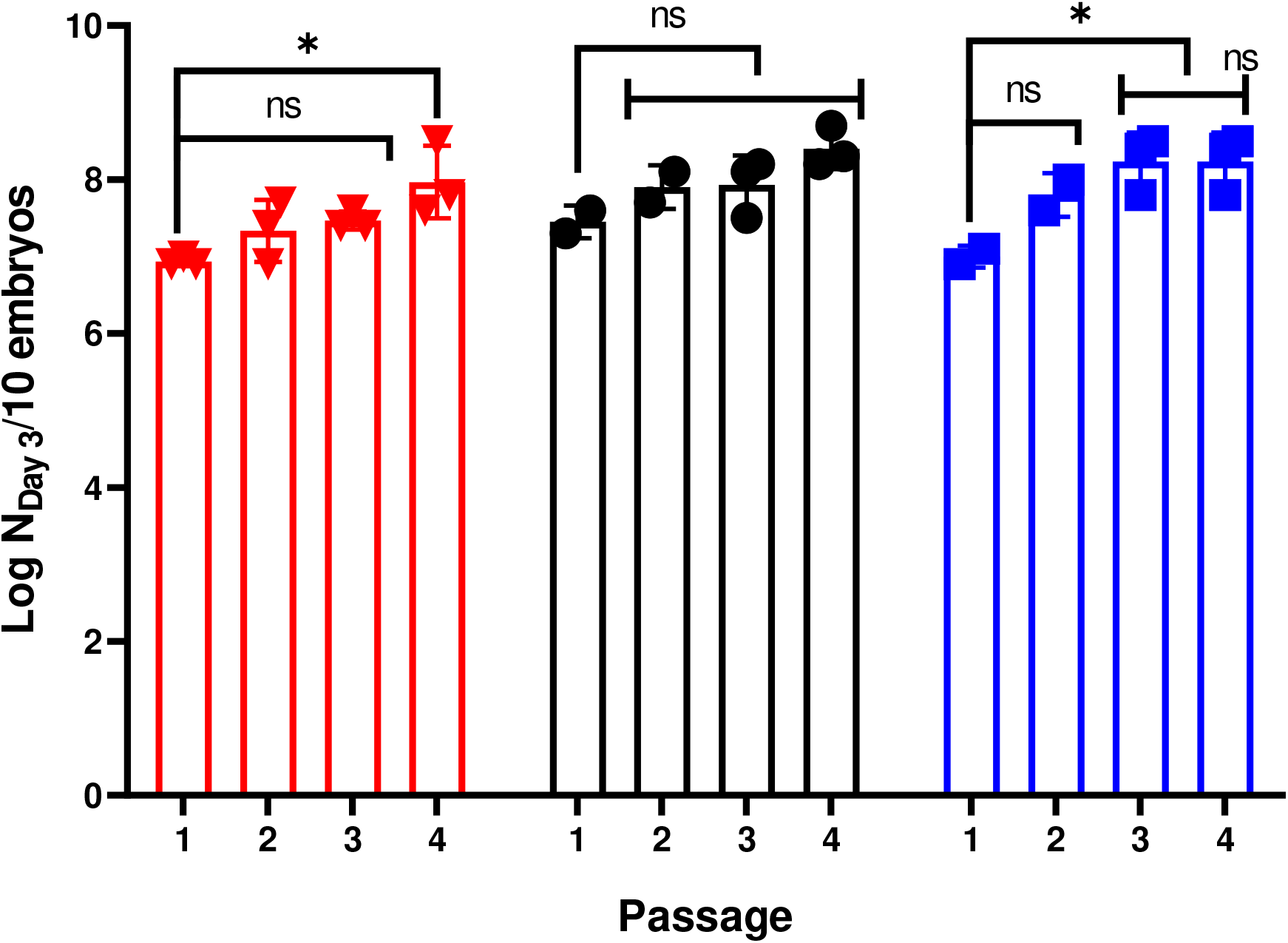
RT-qPCR detection of hNoV GII.4[P16] (red triangles), GII.2[P16] (black circles), and GII.17[P31] (blue squares) injected to zebrafish embryos at 3 dpi of different passages. Each data point was from a pool of 10 embryos. Each column represents the average the independent biological samples, and each error bar indicates the standard deviations. *, significantly different with P < 0.05; ns, no significance with P > 0.05.

### The zebrafish embryo tool enabled direct profiling of hNoV binding to human histo-blood group antigens (HBGAs)

The pathogenesis of hNoVs have been demonstrated to be closely correlated with HBGAs (22, 23). HBGAs are a group of complex terminal carbohydrates can be found as soluble antigens in saliva and are expressed on the mucosal epithelium of the digestive tract (24). Therefore, human clinical samples (stool samples) are typically not used to study the binding between the viruses and HBGAs, largely due to the presence of various types and abundance of HBGA in the stool samples. Preparation of baculovirus-expressed virus-like particles (VLPs), on the other hand, is with high cost and technical issues including the non-uniformity in size during assembly and the particle dissociation during storage may happen as reported previously (25). In addition, gene cloning and baculovirus-expression might introduce certain discrepancies in between of genuine virus particles and VLPs. Here, as the hNoVs produced by zebrafish embryos are with both high titers and clean background, the binding patterns of the three tested hNoV strains were characterized by a saliva-binding RT-qPCR to compare the stool samples and zebrafish embryo amplified samples (passage 4).

Four saliva samples from different individuals with representative HBGA phenotypes were used, including two secretors (S1 and S2) and two non-secretors (NS1 and NS2) based on the detection of Fucα1-2Gal-R. According to the ABO(H) typing, S1 was type A and S2 was type B. Both S1 and S2 were positive for Lewis b and y antigens and negative for Lewis a and x antigens. For non-secretors’ saliva, both NS1 and NS2 were negative for all of the ABO(H) antigens. NS1 was positive for all four Lewis antigens and NS2 was only positive for Lewis a antigen (Table 1).

**Table 1.**
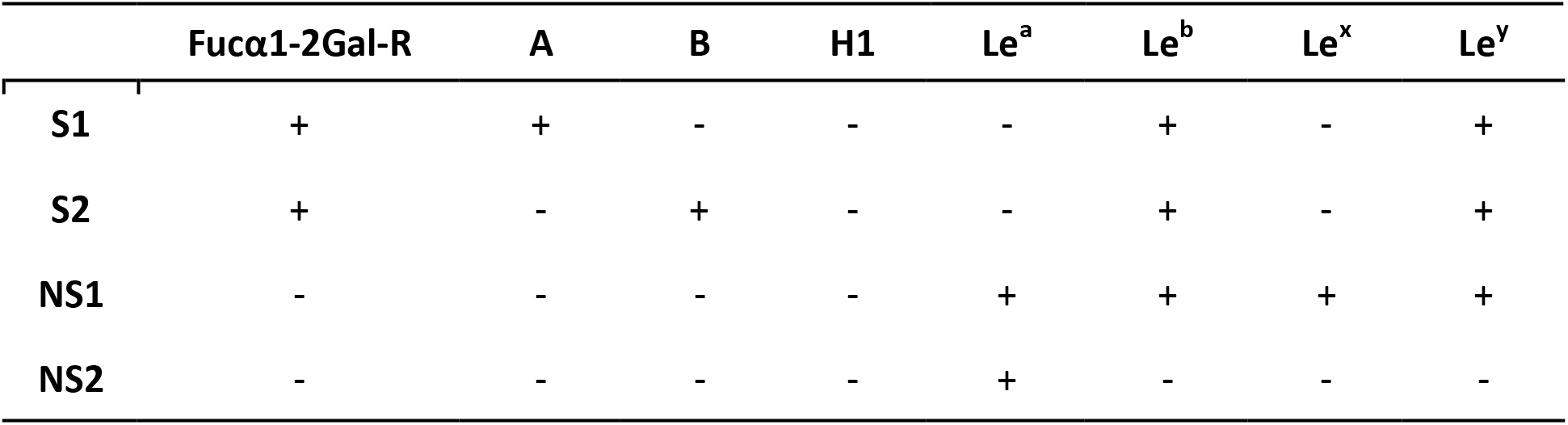
Phenotypes of the saliva samples used in this study.

The binding levels of the hNoVs and HBGAs in saliva were characterized by a “binding indicator” defined as the logarithms of the bound virus ratios (binding indicator = log [genome copies of hNoVs bound on saliva coated plates / genome copy numbers of hNoVs in the samples added to the plates]) as recorded in Figure 3. As expected, the hNoVs amplified by zebrafish embryos indeed showed clearer binding patterns than the viruses in stool samples. For all of the three strains tested in this study, hNoVs GII.4[P16], GII.2[P16], and GII.17[P31], viruses from zebrafish embryos showed significantly higher binding levels than viruses from stool samples to the secretors’ saliva S1 and S2 (P < 0.05), while no such significance was observed in the binding to the non-secretors’ saliva NS1 and NS2 (P > 0.05), except GII.4[P16] to NS1 (P < 0.05). In the meantime, the binding levels of all three strains from zebrafish embryos to the secretors’ saliva S1 and S2 were also significantly higher than to the non-secretors’ saliva NS1 and NS2 (P < 0.05). These results were in great accordance with the clinical records reported by researchers from multiple countries and regions in the world, that the non-secretor individuals, who only express Lewis antigens, are less susceptible to hNoV infection (22). Therefore, this hNoV replication tool with the use of zebrafish embryos supplied an efficient and straightforward way to investigate the binding patterns of the hNoV strains to HBGA without the need of preparing VLPs.

**Figure 3.**
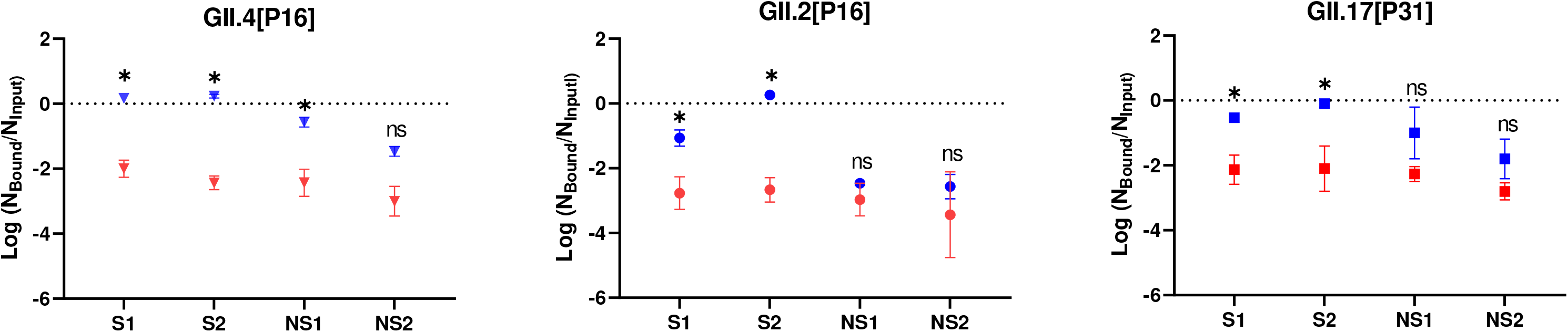
Saliva-binding RT-qPCR results of hNoV GII.4[P16] (triangles), GII.2[P16] (circles), and GII.17[P31] (squares) from stool samples (red triangles, circles and squares) and from zebrafish embryos, passage 4 (blue triangles, circles and squares). The y-axis demonstrated the virus binding levels (binding indicator = log [genome copies of viruses bound on saliva coated plates / genome copy numbers in the samples added to the plates]). Each data point represents the average of triplicates, and each error bar indicates the standard deviations. *, significantly different with P < 0.05; ns, no significance with P > 0.05.

### The zebrafish embryo tool enabled evaluation of hNoV inactivation identifying up to 2-log reduction

For viruses well-adapted to cell culture, the virus infectivity is usually titered by plaque assay or 50 % cell culture infectious dose (TCID_50_) endpoint dilution assay, of which both remain challenging for hNoVs. The current reports evaluating hNoV inactivation efficacy with the use of HIEs were mainly for (qualitative) validation purposes, confirming a certain treatment could inactivate the viruses to non-infectious levels (26–28). Previously, with the use of zebrafish larvae, we managed to titer hNoV as TCID_50_ units using the Reed and Muench calculation method (29). However, only 0.8 log virus reduction by heat treatment was able to be identified, not only due to the limited virus titer in the stool samples but also because the stool samples were with complex matrix and thus needed to be diluted to minimize the matrix effect (19). The zebrafish embryos, on the other hand, generated viruses with higher titers and clean background by continues passaging and thus enabled a larger scale of log reductions to be identified.

In this study, a dilution series of the three hNoV strains GII.4[P16], GII.2[P16], and GII.17[P31] produced by zebrafish embryos (passage 4, ~ 8 log genome copies/10 embryos lysed in 100 μL) were first tested in parallel. Since each data point was from a pool of 10 zebrafish embryos, when the viruses were 10 and 100 times diluted, the virus replication showed large variations in between of the biological replicates. All three tested virus strains became non-infectious consistently when they were 1000 time diluted (Figure 4a). Thus when the same samples (produced by zebrafish embryos, passage 4, ~ 8 log genome copies/10 embryos lysed in 100 μL) were inactivated and showed consistently no infectivity after the treatment, we were able to make a conservative estimation that the treatment was able to inactivate the hNoVs by at least 2 logs. In comparison with our previous study (19) which was very laborious and each data point was generated from 240 zebrafish larvae, this system enabled a more efficient evaluation of hNoV inactivation.

**Figure 4.**
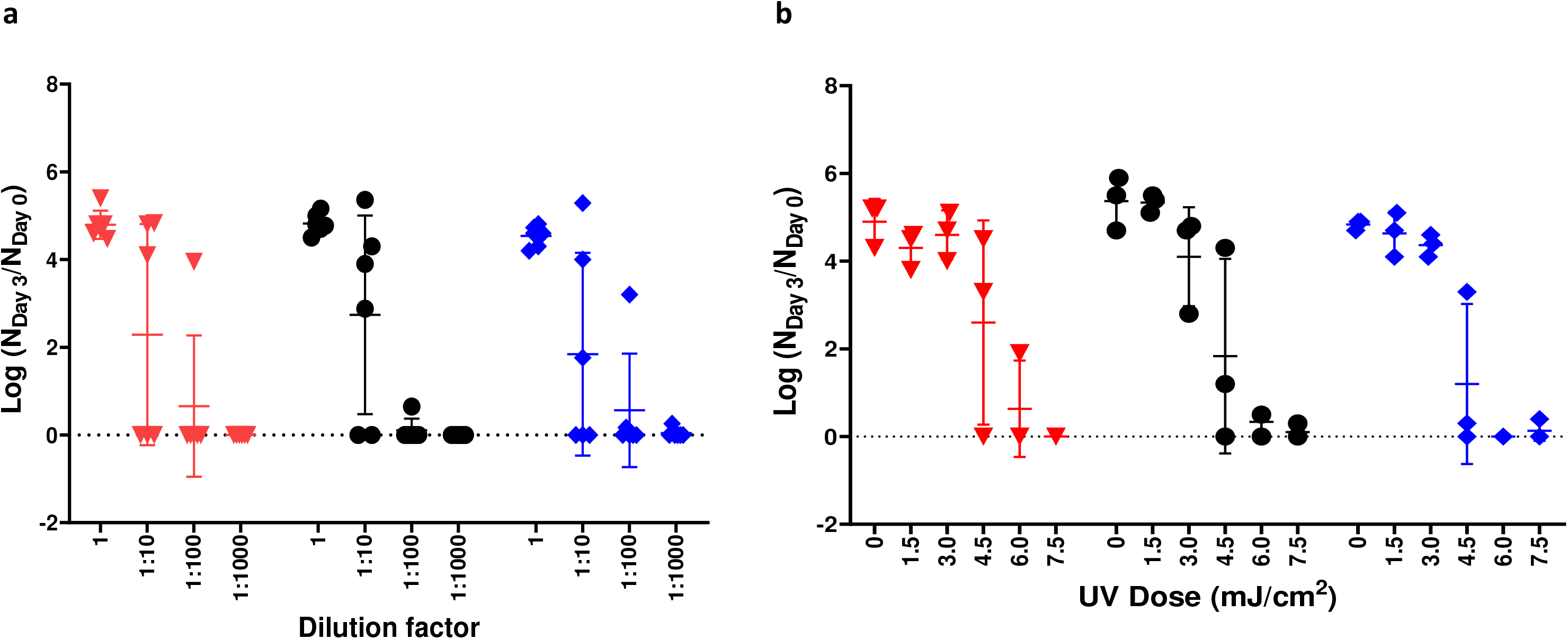
Increase of hNoV injected to zebrafish embryos at 3 dpi in comparison with 0 dpi (log N_Day 3_/N_Day 0_) along with the dilution of virus suspensions (a) and after UV_254_ treatment at different doses (b). Each data point was from a pool of 10 embryos. Red triangles represent data of hNoV GII.4[P16], black circles represent data of GII.2[P16], and blue squares represent data of GII.17[P31]. Each column represents the average the independent biological samples, and each error bar indicates the standard deviations.

In order to demonstrate this concept, in the following, we measured the inactivation of the three hNoV strains with a series of UV irradiation doses at 254 nm (UV_254_, from 1.5 mJ/cm^2^ to 7.5 mJ/cm^2^) and showed that a dose of 6 mJ/cm^2^ UV_254_ was able induce > 2-log hNoV infectivity reduction for all three strains tested (Figure 2b). UV_254_ is a commonly used disinfection process in water treatment as well as for environmental surfaces including food processing facilities. Researchers have been dedicating numerous efforts in evaluating the efficacy of UV on enteric viruses till very recent years (30, 31). In 2020, Rockey et al. (31) reported a genome-wide PCR-based approach evaluating UV disinfection of hNoVs and constructed a linear inactivation curve with an inactivation constant of 0.27 cm^2^/mJ (thus 1.62-log reduction after 6 mJ/cm^2^ UV_254_ treatment). As discussed by the authors, this result was a significant departure from previous reports of 0- to 1.5-log inactivation after UV_254_ doses as high as 300 mJ/cm^2^, which relied on short regions of the genome (32–34). As a further improvement, in this study, by measuring the virus infectivity directly, we revealed that hNoVs were not as UV resistant as previously predicted by PCR-based methods (> 2-log reduction after 6 mJ/cm^2^ UV_254_ treatment for hNoV GII.4[P16], GII.2[P16], and GII.17[P31]). This result also suggests that hNoVs are generally more UV susceptible than multiple enteric viruses and commonly used hNoV surrogates as tested before (0.84-log reduction for bacteriophage MS2, 1.86-log reduction for Echovirus 12 [EV12], 1.92-log reduction for murine norovirus [MNV] [31], 1.08-log reduction for Tulane virus [TV], 0.50-log reduction for rotavirus [RV] [30] after 6 mJ/cm^2^ UV_254_ treatment, calculated based on the linear inactivation curves as reported).

## CONCLUSIONS

In summary, the zebrafish embryo tool as reported in this study represents an essential improvement in hNoVs replication method. With high efficiency and robustness, this tool was able to strengthen hNoV related research in a versatile way. In pathogenesis study, the zebrafish embryo generated viruses were with clear binding patterns to HBGAs in human saliva. In disinfection study, this tool was used to show that a dose of 6 mJ/cm^2^ UV254 was able induce > 2-log hNoV infectivity reduction for all three strains tested, suggesting that hNoVs were more UV susceptible than multiple enteric viruses and commonly used hNoV surrogates as tested before. In the future, it is expected that this tool will not only benefit epidemiological research on hNoV but also be used to generate hNoV inactivation parameters which are highly needed by the water treatment and food industry.

## MATERIALS AND METHODS

### HNoVs from stool samples

HNoVs in stool samples were kindly provided by the Molecular Laboratory, Department of Molecular Pathology, Singapore General Hospital and genotyped in our laboratory at National University of Singapore over the B-C genome region (35). In order to prepare the virus suspension for injection, an aliquot of 100 mg of the stool sample was suspended in 1 mL of sterile phosphate-buffered saline (PBS), thoroughly vortex and centrifuged for 5 min at 9,000 × *g*; the supernatant was harvested and stored at - 80 °C until further use.

### Zebrafish maintenance and husbandry

Wild type adult zebrafish (*Danio rerio*) were maintained in the aquatic facility of National University of Singapore with water temperature at 28 °C and 14/10 h light/dark cycle. Fertilized eggs were collected from adults placed in mating cages and kept in petri dishes containing embryo medium E3 (5 mM of NaCl, 0.17 mM KCl, 0.33 mM of CaCl_2_, 0.33 mM MgSO_4_). All zebrafish experiments were performed in compliance with the Institutional Animal Care and Use Committee guidelines, National University of Singapore.

### Microinjection of zebrafish larvae and embryos in the yolk

Seventy-two hour post-fertilised larvae were first anaesthetized for 2 - 3 min in embryo medium E3 containing 0.2 mg/mL Tricaine (Sigma-Aldrich, USA). After which, they were aligned on the agarose surface with their yolk facing the microinjection needle. Three nL of the viral suspension was injected into the yolk of each larva. For embryo injection, fertilised eggs were collected and washed with embryo medium E3 three times. Prior to injection, embryos were inspected under an inverted microscope to ensure that the embryos had not developed past the spherical stage (~4 hpf). Thereafter, the embryos were transferred to a petri dish with grooves of a mold imprint (6 rows of V-shaped grooves) in 1.5 % agarose. Three nL of the viral suspension was injected into the yolk of each embryo with a pulled borosilicate glass capillary needle. After the injection, the zebrafish (either the embryos or larvae) were transferred to a petri dish containing E3 and further maintained at 29.5 °C. Every day post-infection, the embryos or larvae were observed to ensure that they were properly developing; dead embryos or larvae were removed, and the medium was changed daily. Videos demonstrating the injection procedures were filmed using 0.05 % phenol red (Sigma-Aldrich, USA) solution to simulate the virus suspensions with Nikon motorized fluorescence stereomicroscope (SMZ25-FL) equipped with a Nikon digital camera DS-10.

### Harvesting of zebrafish larvae/embryos for analysis

The zebrafish larvae/embryos were harvested with 10 larvae/embryos pooled as one sample. After being euthanized on ice, the larvae/embryos were mixed with 100 μL PBS in 1.5 mL tubes and homogenized with 3 cycles of 15 s at 6,500 rpm with rest intervals of 60 s by FastPrepTM 24-5G tissue-cell homogenizer (MP Biomedicals, USA). The homogenates were clarified by centrifugation at 9,000 *× g* for 5 min and the supernatants were used as the virus suspension for RT-qPCR analysis or further investigation (passaging of viruses by being injected to new zebrafish embryos, profiling of binding capabilities to HBGAs, UV inactivation study).

### RNA extraction and RT-qPCR for hNoV detection

RNA was extracted using RNeasy Mini Kit (Qiagen, Germany) following the manufacturer’s protocol.

RT-qPCR analyses of NoV GII were carried out using GoTaq^®^ Probe 1-Step RT-qPCR System (Promega, USA). Primers and the FAM/TAMRA-labelled probe of hNoV GII were according to ISO 15216-1:2017. Forward primer QNIF2: 5’-ATGTTCAGRTGGATGAGRTTCTCWGA-3’; reverse primer G2SKR: 5’-TCGACGCCATCTTCATTCACA-3’; probe QNIFs: 5’ FAM-AGCACGTGGGAGGGCGATCG-3’ TAMRA. Cycling conditions were 45°C for 15 min, then 95°C for 10 min, followed by 40 cycles with 95 °C for 15 s and 60 °C for 30 s in each cycle. Cycle threshold (Ct) values were determined during RT-qPCR analysis using StepOneTM system (Applied Biosystems, USA).

Double-stranded DNA (dsDNA) containing the specific primers-probe binding sites were synthesized for NoV GII and cloned into the pGEM-T Vector (Promega), resulting in the NoV-GII plasmids. The plasmids with inserts were purified by using a Plasmid Midi Kit (Qiagen). Plasmid concentration was determined by photospectroscopy at 260 nm using the BioDrop DuoTM spectrophotometer (BioDrop, United Kingdom). Ten-fold serial dilutions ranging from 5 × 10^6^ to 5 copies of all positive control plasmids were used to prepare a standard curve and enumerate the NoV GII.

### Detection of Fucα1-2Gal and HBGA phenotyping in saliva samples

Human saliva samples were collected from healthy adults under the ethical approval of National University of Singapore Institutional Review Board (NUS-IRB, reference code: N-20-035).

The saliva samples were boiled at 95 °C for 10 min, followed by centrifugation at 10, 000 *x g* for 5 min. The supernatant was collected and diluted at 1:100 by carbonate: bicarbonate buffer (pH 9.6; CBS) and was used to coat 96-well microtiter plate (100 μL/well) at 4 °C overnight. The coated plate was washed three times with PBS containing 0.05 % Tween 20 (PBS-T) and blocked with 5 % nonfat dried milk (Blotto, 200 μL/well) at 37 °C for 1 h. In order to detect Fucα1-2Gal-R, which is specifically present in secretor but not in non-secretor saliva, HRP-conjugated Ulex europaeus agglutinin (UEA-I, Sigma-Aldrich, USA) diluted at 1:3200 (starting conc. 1 mg/mL) was added and incubated for 1 h at 37 °C. After three washes with PBS-T, the presence of Fucα1-2Gal-R was detected with a TMB (3,3’,5,5’-tetramethylbenzidine) kit (Sigma-Aldrich, USA), and the signal intensities [the optical density at 450 nm (OD_450_)] were read with a Multiskan Sky plate reader (Thermo Fisher Scientific, China). Similarly, for the determination of the HBGA phenotype of the saliva samples, the blocked wells were added with 1:200 diluted HBGA monoclonal antibody anti-A (BG 2), anti-B (BG 3), anti-H type 1 (BG 4), anti-Lewis a (BG 5), anti-Lewis b (BG 6), anti-Lewis x (BG 7), and anti-Lewis y (BG 8) (MAb; Covance, Emeryville, CA, USA) and then incubated for 1 h at 37 °C. After three washes with PBS-T, the HBGA phenotype was detected with horseradish peroxidase conjugated with goat anti-mouse IgG (1:1500; Sigma-Aldrich, USA) for type A, H, and Lewis a, and goat anti-mouse IgM (1:1500; Sigma-Aldrich, USA) for type B and Lewis b, x and y. Horseradish peroxidase activity was detected with a TMB kit, and the OD_450_ were read with the Multiskan Sky plate reader. The cutoff value was taken as two times the absorbance value of the negative controls as previously performed by Nordgren et al. (2019).

### HNoVs binding to saliva

Microtiter plate was coated with saliva samples at 1:100 with CBS (100 uL/well) at 4 °C overnight. The coated wells were washed three times with PBS-T, blocked with 5 % nonfat dried milk at 37 °C for 1 h. Thereafter, hNoV samples from human stool or zebrafish embryo amplified virus suspensions prepared as described above were added at 10^6^ virus genome copies/well and incubated for 1 h at 37 °C. After three washing with PBS-T, direct lysis was performed by adding lysis buffer of the RNeasy Mini Kit (Qiagen) to individual wells, and they were subsequently subjected to RNA extraction according to the manufacturer’s protocol. RT-qPCR analysis of the bound viruses were performed as described above.

### UV_254_ treatment of viruses

The UV treatment was performed in a biosafety cabinet (ESCO, Singapore, LA2-4A1-E) with low-pressure mercury lamps producing mainly 254-nm UV light (UV-30A). The viral suspensions prepared as described above (zebrafish embryos amplified, passage 4) were spotted into a sterile petri dish (20 μL as one droplet) and exposed to the UV light till reaching a dose of 1.5, 3.0, 4.5, 6.0 and 7.5 mJ/cm^2^ respectively. The UV irradiance values were measured by an optical radiometer (UVX Radiometer, Analytik Jena, USA). After treatment, the samples were recovered by pipetting and homogenizing for several times and proceeded to injection to zebrafish embryos directly.

### Statistical analysis

Statistical analyses were performed using the software SPSS for windows, version 22.0. A paired t-test was used for data with two groups, one-way analysis of variance was used for data with more than two groups. Significant differences were considered when P was < 0.05.

## AUTHOR CONTRIBUTION STATEMENT

MTHT and DL conceived and designed research. MTHT conducted experiments. ZG contributed analytical tools. MTHT and DL analyzed data. MTHT and DL wrote the manuscript. All authors read and approved the manuscript.

## ACKNOWLEDGMENT

This study was supported by a Ministry of Education (MOE) academic research fund (AcRF) project A-0008418-00-00 (PI: Dan Li, Jul 2021 to Jun 2023).

